# Therapeutic Targeting of Microglial Hexokinase-2 Recalibrates Inflammasome Activation and Improves Functional Recovery After Traumatic Brain Injury

**DOI:** 10.64898/2026.03.01.708896

**Authors:** Claudia Mera-Reina, Juan F. Codocedo, Paul B. Fallen, Jack Scott, Cristian A. Lasagna-Reeves, Gary E. Landreth

## Abstract

Traumatic brain injury (TBI) initiates a secondary inflammatory cascade in which sustained microglial activation contributes to long-term neurological dysfunction. Microglial inflammatory states depend on glycolytic reprogramming, suggesting that targeted modulation of metabolic regulators may attenuate post-traumatic inflammation while preserving essential immune functions.

Hexokinase-2 (HK2), a rate-limiting glycolytic enzyme, regulates inflammatory signaling and inflammasome activation in microglia in neurodegenerative contexts; however, its role in TBI remains undefined. We therefore examined whether partial suppression of microglial HK2 modulates inflammatory responses following severe TBI.

HK2 was robustly induced in microglia during the sub-acute phase after injury. Pharmacological inhibition of HK2 improved motor coordination without impairing locomotion or cognitive performance and selectively reduced inflammasome-related gene expression and ASC accumulation, particularly within the hippocampal hilus. Importantly, HK2 antagonism slowed microglial proliferation while preserving efferocytic capacity. Partial genetic reduction of microglial HK2 phenocopied these molecular and behavioral effects, supporting an HK2-dependent mechanism.

Together, these findings identify microglial HK2 as a therapeutically targetable regulator of inflammatory amplification after TBI. Partial modulation of this pathway attenuates secondary neuroinflammation while maintaining critical microglial functions, highlighting HK2 as a promising strategy to improve functional recovery after traumatic brain injury.

## Introduction

Traumatic brain injury (TBI) is a leading cause of death and long-term neurological disability worldwide. Beyond the initial mechanical insult, secondary injury processes, including excitotoxicity, oxidative stress, and neuroinflammation, drive progressive neuronal dysfunction and limit functional recovery [1,2]. Among these secondary mechanisms, neuroinflammation is increasingly recognized as a major determinant of outcome after TBI.

Microglia, the resident immune cells of the central nervous system, are central regulators of the inflammatory response to brain injury. Following TBI, microglia rapidly activate and adopt diverse phenotypes that can promote tissue repair through debris clearance and remodeling but can also exacerbate damage through sustained inflammatory signaling [3,4]. Thus, the balance between protective and maladaptive microglial responses critically shapes the progression of secondary injury. Identifying molecular mechanisms that regulate this balance remains a major therapeutic priority.

Recent work has established that microglial activation is tightly coupled to metabolic reprogramming. In response to injury or neurodegenerative stimuli, microglia shift from oxidative phosphorylation toward glycolysis, a transition that supports proliferation, cytokine production, and phagocytic activity [5–7]. This metabolic shift is not merely a consequence of activation but a prerequisite for the acquisition of inflammatory functions. Accordingly, targeting key regulators of glycolytic flux has emerged as a strategy to modulate microglial immune responses without abolishing essential homeostatic functions [8].

Hexokinase-2 (HK2), the rate-limiting enzyme of glycolysis, has emerged as a regulator of immune cell activation beyond its canonical metabolic function. In microglia, HK2 modulates inflammatory signaling and inflammasome activation, linking glucose utilization to innate immune responses [9]. Although glycolytic reprogramming is well documented in TBI models [6], whether HK2 specifically governs microglial inflammatory responses after traumatic injury remains unexplored. Furthermore, it remains unknown whether therapeutic modulation of HK2 can recalibrate microglial activation without compromising essential repair functions.

Here, we hypothesized that partial therapeutic targeting of microglial HK2 after severe TBI would attenuate inflammasome activation and limit maladaptive inflammatory amplification while preserving fundamental microglial functions, thereby improving functional recovery. Using a controlled cortical impact (CCI) model, we demonstrate that HK2 is robustly upregulated in microglia following injury. Pharmacological antagonism of HK2 improves motor performance and selectively suppresses inflammasome-associated gene expression and ASC accumulation in a region-dependent manner. Importantly, partial genetic reduction of microglial HK2 phenocopies these effects, supporting a specific HK2-dependent mechanism. Together, our findings identify microglial HK2 as a therapeutically targetable regulator of inflammatory tone after TBI and suggest that partial modulation of this pathway may mitigate secondary injury while preserving essential immune functions.

## Results

### HK2 is upregulated in microglia in response to CCI-induced TBI

Considering the reported role of HK2 in regulating microglial activation and inflammatory signaling in neurodegenerative conditions, we investigated whether HK2 expression is similarly induced following severe TBI. To this end, we assessed HK2 protein levels in brain sections from mice subjected to CCI. Immunohistochemical analysis performed 15 days post-injury revealed a marked increase in HK2 immunoreactivity within Iba1⁺ microglia in the ipsilateral cortex adjacent to the injury site compared to the contralateral hemisphere (Figure 1A–B). Elevated HK2 expression was also observed in the ipsilateral hippocampus (Figure 1A–C), indicating that HK2 induction extends beyond the lesion core.

**Figure 1.**
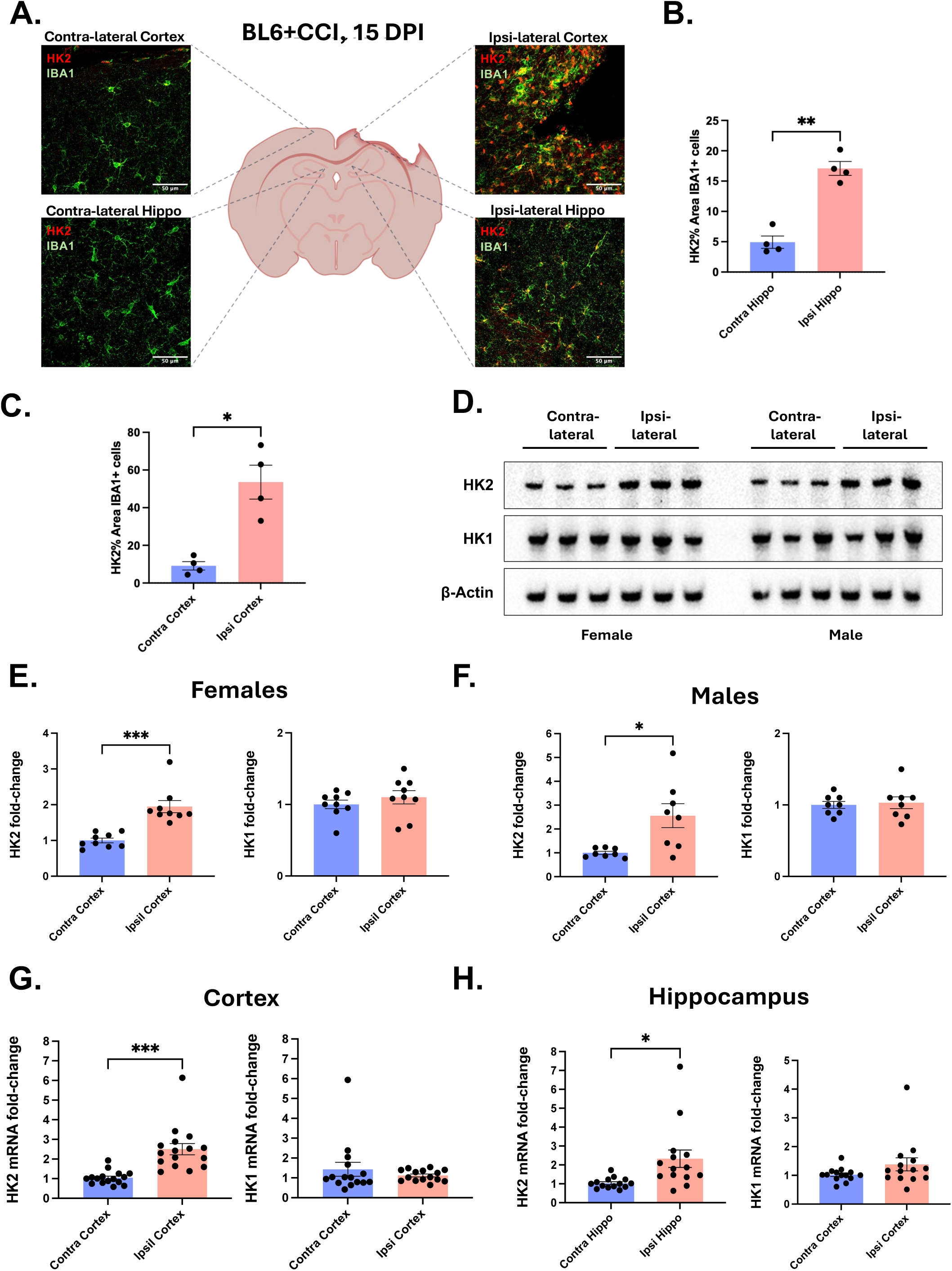
HK2 is transcriptionally and translationally upregulated in microglia following CCI-induced TBI. **(A–C)** Representative immunohistochemical images showing HK2 expression in IBA1⁺ microglia in the ipsilateral and contralateral cortex and hippocampus 15 days after controlled cortical impact (CCI). Increased HK2 immunoreactivity is observed in the ipsilateral cortex adjacent to the injury site and in the ipsilateral hippocampus with respect to its contralateral side. (*n* = 4, *p < 0.05 and **p < 0.01, paired T test). **(D–F)** Western blot analysis and quantification of HK2 and HK1 protein levels in cortical lysates from male and female mice 15 days post-CCI. HK2, but not HK1, is significantly increased following injury in both sexes. (*n* = 9 females and n=8 males, *p < 0.05 and ***p < 0.001, paired T test) **(G–H)** qPCR analysis of HK2 mRNA expression in cortex and hippocampus confirms transcriptional induction after CCI. (*n* = 15 females and n=14 males, *p < 0.05 and ***p < 0.001, paired T test)

To quantify these changes and assess potential sex differences, we performed Western blot analyses on hippocampal lysates from male and female mice 7 days post-CCI. HK2 protein levels were significantly elevated in both sexes, with no detectable difference between males and females (Figure 1E–F). In contrast, HK1, the predominant neuronal hexokinase isoform, remained unchanged following injury (Figure 1E–F), supporting the specificity of HK2 induction.

Consistent with these findings, qPCR analysis of cortical and hippocampal tissue confirmed transcriptional upregulation of HK2 (Figure 1G–H), indicating that HK2 induction is part of the injury-associated gene expression program.

Together, these results demonstrate that severe TBI robustly and selectively induces microglial HK2 across both cortical and hippocampal regions. The regional breadth and isoform specificity of this response suggest that HK2 participates in secondary inflammatory signaling after injury.

### HK2 inhibition improves motor coordination after TBI without affecting anxiety-like behavior or locomotion

Having established that HK2 is robustly induced following CCI, we next asked whether early pharmacological targeting of HK2 could modify functional outcomes after severe TBI. Because excessive microglial inflammatory amplification contributes to secondary injury progression, we tested whether transient HK2 antagonism during the acute phase would influence behavioral recovery. Mice received the HK2 antagonist lonidamine (LND; 50 mg/kg, i.p.) beginning 24 hours after CCI for 7 consecutive days. Behavioral testing was conducted at 15 days post-injury using open field, Y-maze, and rotarod paradigms (Figure 2A).

**Figure 2.**
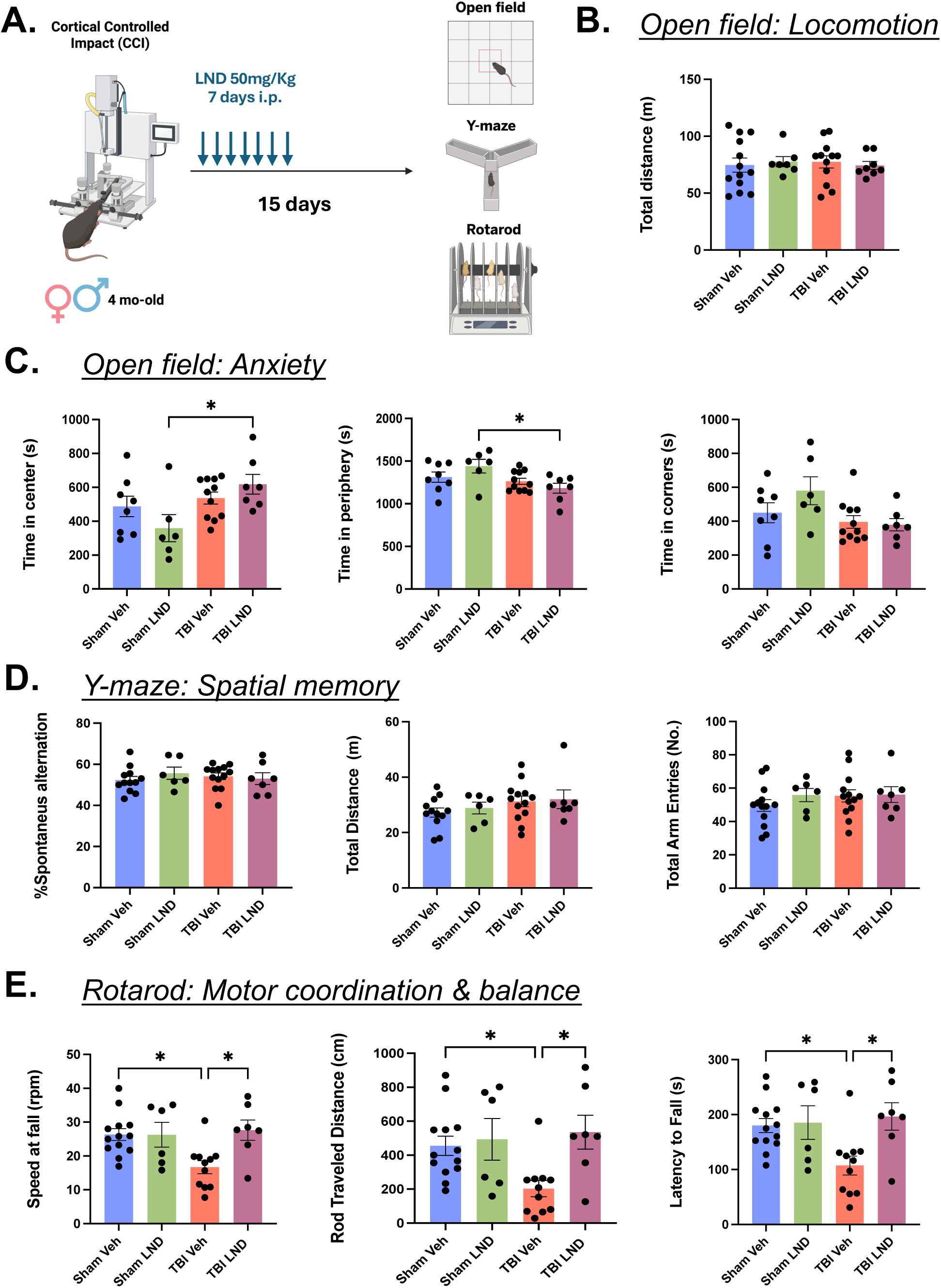
Pharmacological HK2 antagonism improves motor coordination after TBI without altering locomotion, anxiety-like behavior, or spatial memory. **(A)** Experimental timeline illustrating CCI, lonidamine (LND) treatment, and behavioral testing. **(B)** Open field total distance traveled shows no differences in locomotor activity across groups. **(C)** Open field center exploration indicates a modest effect in center time following LND treatment in the Sham mice group without evidence of injury-induced anxiety-like behavior. (*n* = 7-13 per group, *p < 0.05, Ordinary one-way ANOVA with post hoc Tukey’s multiple comparisons) **(D)** Y-maze spontaneous alternation reveals no differences in short-term spatial working memory among groups. Likewise, no changes were observed in locomotion and exploratory behaviors. **(E)** Rotarod performance demonstrates significant impairment after CCI, which is completely rescued by LND treatment, as shown by increased latency to fall, distance traveled, and maximal speed. (*n* = 7-13 per group, *p < 0.05, Ordinary one-way ANOVA with post hoc Tukey’s multiple comparisons)

Open field analysis revealed no differences in total distance traveled among groups, indicating that neither CCI nor LND altered baseline locomotion (Figure 2B). Time spent in the center was modestly increased by LND treatment without changes in overall activity (Figure 2C), suggesting a subtle anxiolytic-like effect rather than injury-related behavioral disruption. No group exhibited pronounced anxiety phenotypes, minimizing confounding effects on motor performance.

Short-term spatial working memory assessed by the Y-maze showed no differences in spontaneous alternation or exploratory behavior across groups (Figure 2D), indicating preserved cognitive function at this post-injury time point.

In contrast, significant differences emerged in the rotarod test (Figure 2E). Mice subjected to severe TBI displayed impaired maximal speed, distance traveled, and latency to fall compared to sham controls. Notably, LND treatment significantly improved all three parameters in injured mice, indicating partial rescue of motor coordination and endurance. These effects were absent in sham animals, suggesting that HK2 inhibition selectively ameliorates injury-induced motor dysfunction.

Sex-segregated analyses (Supplementary Figure 1) showed that motor improvement with LND was primarily observed in males, whereas females did not exhibit significant changes. Although not powered to formally assess sex differences, this pattern parallels previously reported male-biased responsiveness to LND in neurodegenerative models, suggesting that pharmacological targeting of HK2 may engage sex-dependent inflammatory mechanisms.

Exploratory analyses of synaptic proteins revealed no change in PSD-95 levels, whereas synaptophysin expression, reduced after TBI, was partially restored by LND in the male cortex (Supplementary Figure 1). While correlative, these findings align with behavioral improvement and suggest possible preservation of presynaptic integrity.

Collectively, these findings reveal a selective behavioral profile following early HK2 antagonism. Despite the severity of cortical injury, transient inhibition of HK2 during the acute phase improved motor coordination without impairing locomotion, anxiety-like behavior, or spatial working memory. The preservation of cognitive performance suggests that HK2 targeting does not globally suppress neural or immune function but rather modulates injury-specific dysfunction. The male-biased responsiveness further highlights a pharmacologically sensitive dimension of inflammatory regulation that warrants deeper mechanistic and pharmacokinetic investigation. Together, these data support the concept that early therapeutic targeting of microglial HK2 may represent a safe and functionally beneficial strategy to improve motor outcomes after severe TBI.

Given that HK2 antagonism improved motor recovery without disrupting baseline behaviors, we next examined how HK2 inhibition reshapes microglial inflammatory programs and whether specific cellular mechanisms underlie these functional benefits.

### HK2 inhibition slows microglial proliferation, does not impair efferocytosis, and attenuates inflammasome activation in a region- and sex-dependent manner

To define the cellular mechanisms underlying the functional benefits of HK2 antagonism, we examined microglial proliferation, debris clearance, and inflammatory gene expression following LND treatment. Using live-cell imaging, we assessed efferocytosis in BV2 microglia co-cultured with fluorescently labeled apoptotic neuroblastoma cells. LND treatment did not alter the efficiency or kinetics of apoptotic cell uptake (Figure 3A,C), indicating that HK2 inhibition does not compromise microglial efferocytosis in vitro. In contrast, continuous confluency monitoring revealed that LND significantly slowed microglial proliferation (Figure 3B), suggesting that HK2 antagonism modulates growth dynamics while preserving essential phagocytic function.

**Figure 3.**
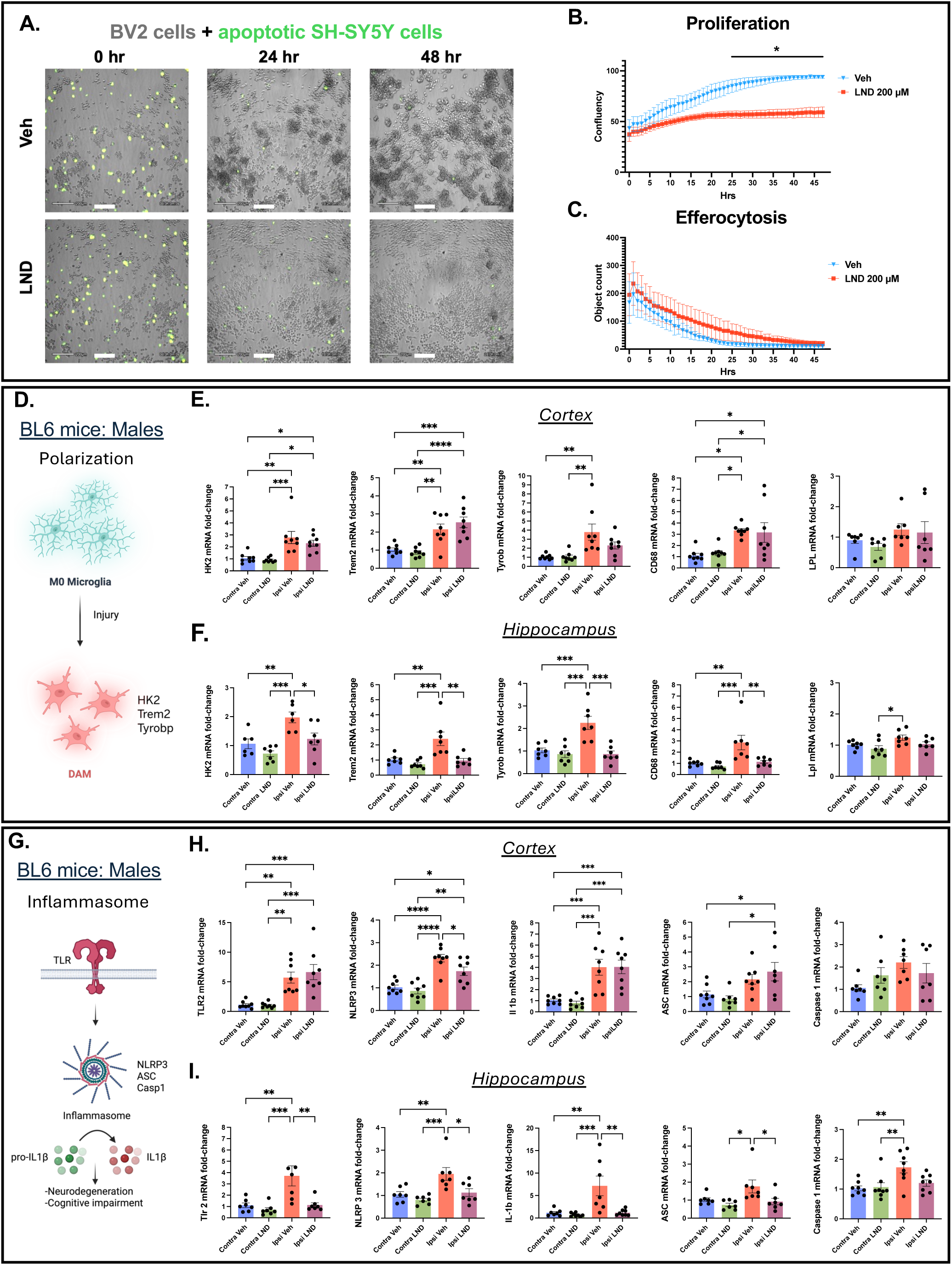
HK2 inhibition slows microglial proliferation without impairing efferocytosis and attenuates microglial gene expression after TBI. **(A)** Live-cell imaging–based efferocytosis assay using BV2 microglia incubated with fluorescently labeled apoptotic SH-SY5Y cells and 200 μM of LND for 48 hrs. **(B)** Quantification of BV2 cell confluency demonstrates reduced proliferation following LND treatment. (*n* =4 per group, *p < 0.05, Multiple unpaired t-test followed by Two-stage linear step-up procedure of Benjamini, Krieger and Yekutieli) **(C)** LND treatment does not alter the efficiency or kinetics of apoptotic cell clearance. **(D–F)** qPCR analysis of a set of microglial activation–associated genes (polarization: HK2, Trem2, Tyrobp, and LPL) in the cortex and hippocampus of male mice following CCI and LND treatment. LND selectively reduces gene induction in the hippocampus. (*n* = 7-8 per group, *p < 0.05, **p < 0.01, and ***p < 0.001, Ordinary one-way ANOVA with post hoc Tukey’s multiple comparisons) **(G–I)** Expression of inflammasome-related genes (Nlrp3, Asc, Casp1, Il1b and Tlr2) is robustly induced after CCI and significantly attenuated by LND in the hippocampus but not cortex. (*n* = 7-8 per group, *p < 0.05, **p < 0.01, and ***p < 0.001, Ordinary one-way ANOVA with post hoc Tukey’s multiple comparisons)

We next assessed microglial transcriptional responses in vivo following CCI. Expression of activation-associated genes, including Trem2, Tyrobp (Dap12), Cd68, LPL, and HK2, was robustly induced in male mice in both injured cortex and hippocampus. LND treatment did not significantly alter gene induction in the cortex but selectively reduced Trem2, Tyrobp, Cd68, and HK2 expression in the hippocampus (Figure 3D–F), indicating region-restricted modulation of microglial activation. HK1 expression remained unchanged across conditions (Supplementary Figure 2A–B), supporting isoform specificity. Notably, LPL levels were not altered at this time point, despite prior reports of LPL upregulation following HK2 antagonism in AD models [10].

We then evaluated inflammasome-related genes (Nlrp3, Asc/Pycard, Casp1, Il1b). TBI induced a strong inflammatory transcriptional response in the cortex that was not modified by LND (Figure 3G–I). In contrast, hippocampal inflammasome components and Il1b were markedly suppressed by LND in male mice, demonstrating selective attenuation of inflammatory amplification in this region.

These transcriptional effects were strongly sex dependent. In female mice, LND did not significantly alter polarization-associated or inflammasome-related gene expression in either cortex or hippocampus (Supplementary Figure 2B–H). This sex-specific regulation parallels the behavioral findings in Figure 2 and is consistent with previously reported male-biased anti-inflammatory responses to LND in 5xFAD AD models [11].

Together, these data indicate that HK2 inhibition selectively limits microglial proliferation and dampens inflammasome activation without impairing efferocytosis. The consistency across distinct models of neurodegeneration and injury reinforces the idea that the proximity to the insult and sex shape microglial metabolic responses to LND. This anti-inflammatory action, particularly evident in the male hippocampus, may contribute to the behavioral recovery observed in LND-treated mice after severe TBI (Figure 2).

### HK2 antagonism attenuates microglial ASC accumulation after TBI

To determine whether the hippocampal suppression of inflammasome-related transcripts was reflected at the protein level, we examined ASC accumulation within IBA1⁺ microglia in cortex and hippocampus following CCI (Figure 4).

**Figure 4.**
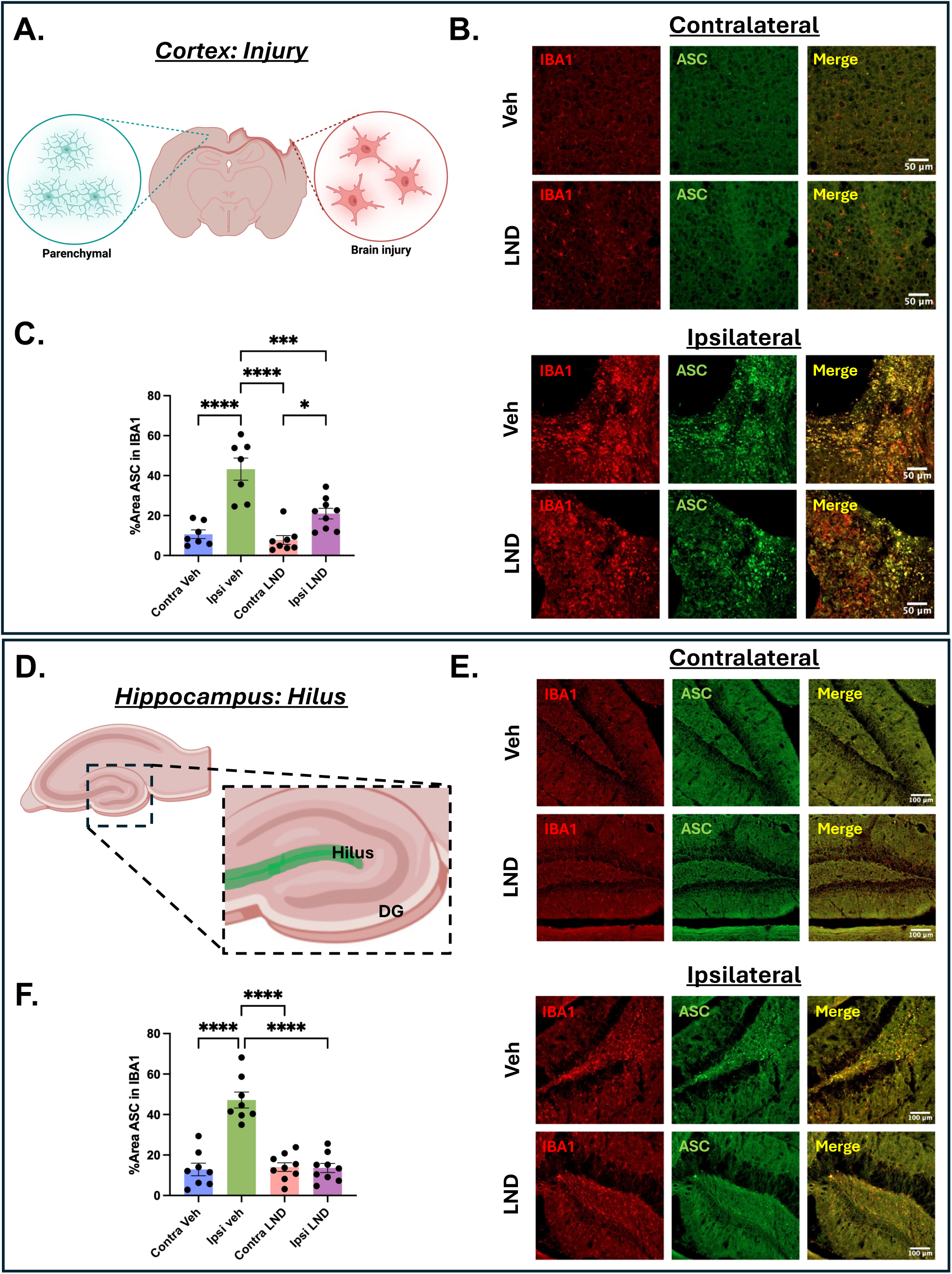
HK2 antagonism reduces microglial ASC accumulation in the injured cortex and hippocampal hilus after TBI. **(A–B)** Schematic and representative immunofluorescence images of IBA1 and ASC in the ipsilateral injured cortex and contralateral cortex. **(C)** Quantification of ASC immunoreactivity normalized to IBA1⁺ area reveals increased inflammasome-associated protein accumulation after CCI that is reduced by LND treatment. (*n* = 7-8 per group, *p < 0.05, ***p < 0.001, and ****p < 0.0001, Ordinary one-way ANOVA with post hoc Tukey’s multiple comparisons) **(D–E)** Schematic and representative images of the hippocampal dentate gyrus highlighting ASC accumulation in IBA1⁺ microglia within the ipsilateral hilus. **(F)** Quantification shows robust ASC induction in the ipsilateral hilus following CCI, which is markedly reduced by LND treatment to levels comparable to contralateral controls. (*n* = 7-8 per group, ****p < 0.0001, Ordinary one-way ANOVA with post hoc Tukey’s multiple comparisons)

CCI markedly increased ASC immunoreactivity within IBA1⁺ microglia in the ipsilateral cortex, consistent with enhanced inflammasome activation at the impact site. Quantification of ASC signal normalized to IBA1⁺ area confirmed increased ASC expression per microglial area rather than changes in cell density. LND treatment significantly reduced ASC immunoreactivity in the injured cortex (Figure 4A–C), demonstrating that HK2 antagonism attenuates inflammasome-associated protein accumulation at the cellular level. Notably, this reduction was detectable despite minimal changes in bulk cortical Asc transcript levels, suggesting that spatially restricted or cell-specific effects may not be captured by region-wide qPCR analysis.

In the hippocampus, ASC induction displayed a strikingly focal distribution. The dentate gyrus hilus is recognized as particularly vulnerable to TBI-induced synaptic remodeling, mossy fiber reorganization, and alterations in neurogenesis, and plays a central role in regulating hippocampal circuit excitability [12–15]. Rather than uniform activation across subfields, increased ASC immunoreactivity was concentrated within IBA1⁺ microglia in the ipsilateral hilus (Figure 4D–F). Quantitative analysis confirmed robust ASC expression in this region following injury. Importantly, LND treatment markedly reduced ASC signal in hilar microglia, restoring levels to those comparable to the contralateral side. The magnitude of suppression was greater in the hippocampus than in cortex, consistent with the region-restricted transcriptional effects observed earlier.

Together, these findings demonstrate that HK2 antagonism limits microglial inflammasome activation in a spatially selective manner, with particularly strong effects in the hippocampal hilus. The normalization of ASC accumulation in this functionally vulnerable region provides a cellular correlate of the regionally dependent inflammatory modulation observed at the transcriptional level and supports a role for HK2 in regulating localized secondary inflammatory signaling after severe TBI.

### Partial genetic reduction of microglial HK2 reproduces the behavioral and molecular effects of pharmacological HK2 inhibition after TBI

To determine whether the effects observed with pharmacological HK2 antagonism were specifically attributable to HK2 regulation in microglia, we employed a tamoxifen-inducible HK2 heterozygous model. As outlined in Figure 5A, HK2^fl/wt mice underwent Cre-mediated recombination in adulthood, followed by CCI at 4 months of age. Behavioral and molecular analyses were performed 15 days post-injury.

**Figure 5.**
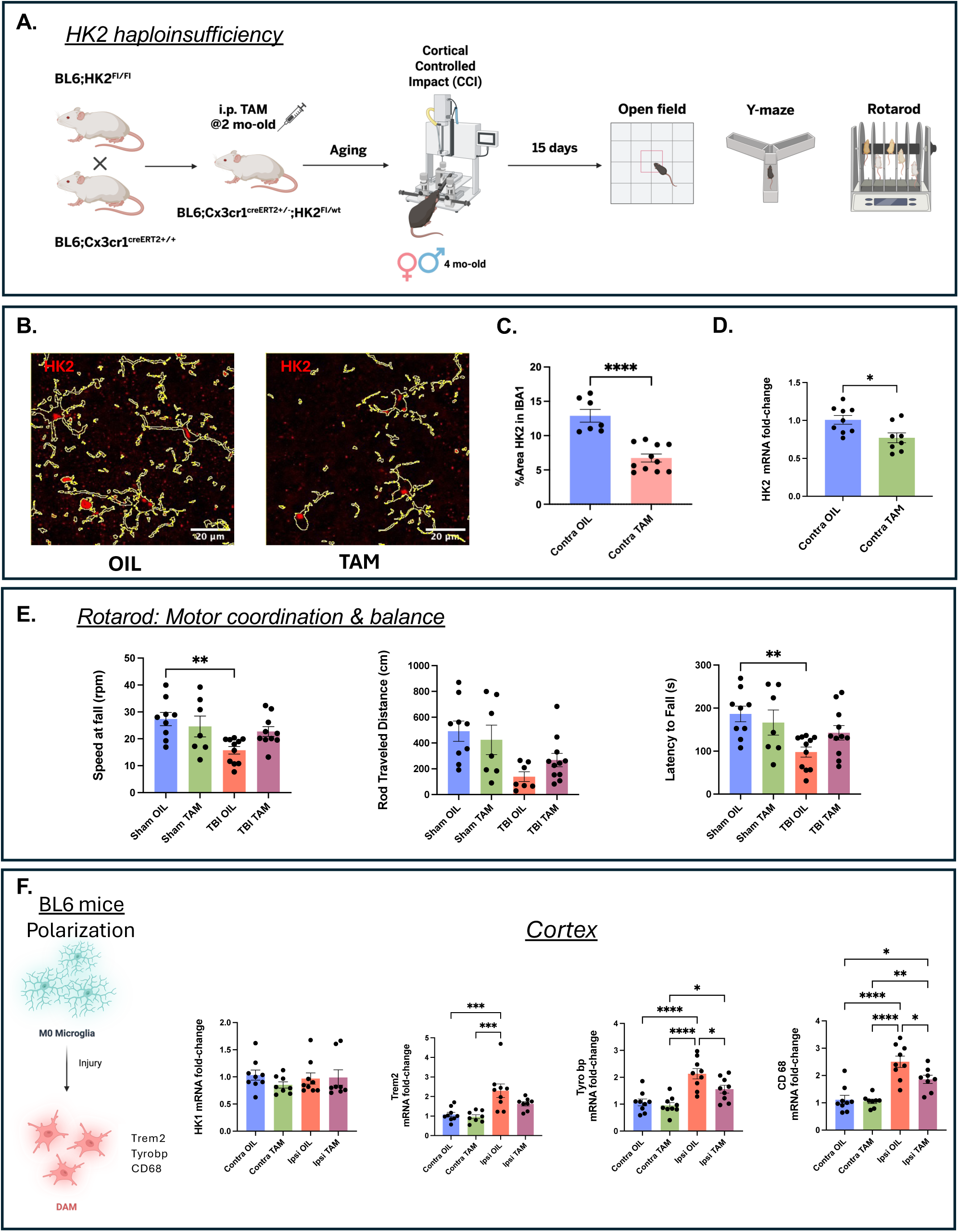
Partial genetic reduction of microglial HK2 reproduces the behavioral and molecular effects of pharmacological HK2 antagonism. **(A)** Schematic of the experimental design using tamoxifen-inducible HK2^fl/wt^ mice, including recombination, CCI, and downstream analyses. **(B–C)** Representative images and quantification showing ∼50% reduction of HK2 immunoreactivity within IBA1⁺ microglia in the contralateral cortex following tamoxifen treatment. (*n* = 7-11 per group, ****p < 0.0001, unpaired T test) **(D)** qPCR validation of reduced HK2 transcript levels in contralateral cortical tissue. (*n* = 8 per group, *p < 0.05, unpaired T test) **(E)** Rotarod performance demonstrating improved motor coordination and endurance in HK2 heterozygous mice after CCI. (*n* = 7-8 per group, ****p < 0.0001, Ordinary one-way ANOVA with post hoc Tukey’s multiple comparisons) **(F)** qPCR analysis showing reduced expression of HK1 and microglial activation markers (Trem2, Tyrobp, Cd68) in cortex following partial HK2 deletion. (*n* = 7-8 per group, *p < 0.05, **p < 0.01, ***p < 0.001, and ****p < 0.0001, Ordinary one-way ANOVA with post hoc Tukey’s multiple comparisons)

Immunohistochemical analysis confirmed efficient partial deletion of HK2 in microglia. In the contralateral cortex, HK2 immunoreactivity within IBA1⁺ cells was reduced by approximately 50% in tamoxifen-treated mice compared with oil-treated controls (Figure 5B–C), consistent with heterozygous recombination. This reduction was corroborated at the transcript level by qPCR (Figure 5D), validating effective HK2 dosage reduction.

Behavioral testing demonstrated that partial genetic reduction of HK2 phenocopied the functional improvements observed with LND treatment. Tamoxifen-treated HK2 heterozygous mice exhibited improved rotarod performance following CCI compared with oil-treated controls (Figure 5E). In contrast, open field and Y-maze performance were unchanged (Supplementary Figure 3), indicating selective improvement in motor coordination without effects on locomotion, anxiety-like behavior, or spatial working memory.

At the molecular level, partial HK2 reduction attenuated microglial activation signatures after injury. qPCR analysis revealed significant decreases in Trem2, Tyrobp, and Cd68 expression in tamoxifen-treated mice relative to controls (Figure 5F). HK1 expression remained unchanged, supporting the specificity of HK2-dependent effects.

Notably, in contrast to the male-biased response observed with pharmacological inhibition, partial genetic reduction of HK2 produced comparable behavioral and transcriptional effects in both sexes. This dissociation suggests that the sex-dependent effects of LND are unlikely to reflect intrinsic differences in HK2 biology, but rather sex-specific pharmacokinetic or pharmacodynamic factors.

Together, these findings establish that partial reduction of microglial HK2 is sufficient to attenuate inflammatory activation and improve motor recovery after severe TBI. The concordance between pharmacological antagonism and genetic dosage reduction strengthens the causal link between HK2 activity and injury-associated inflammatory amplification. Importantly, the preservation of baseline behaviors and essential microglial functions across both approaches supports the view that HK2 functions as a modulable inflammatory checkpoint rather than a global regulator of microglial viability. These results position microglial HK2 as a therapeutically targetable node capable of recalibrating secondary inflammatory responses after brain injury.

## Discussion

TBI remains a major clinical challenge in part because secondary inflammatory processes evolve over days to weeks and contribute substantially to functional decline. Microglia are central regulators of this response, balancing debris clearance and tissue remodeling against sustained inflammatory signaling that exacerbates neuronal dysfunction [4,16]. Strategies that recalibrate microglial activation without impairing repair functions are therefore of high therapeutic interest.

Metabolic reprogramming is a key determinant of microglial activation [17]. Following CNS injury, microglia shift toward glycolysis to support proliferation and cytokine production. HK2 acts as a regulatory node linking glycolytic flux to inflammatory signaling [9], and studies in AD [10,11,18] and stroke [19–21] demonstrate that HK2 enrichment in activated microglia is required for pro-inflammatory phenotypes. Extending this framework to severe TBI, we show now that HK2 is robustly induced not only at the cortical impact site but also in anatomically distant regions, such as the hippocampus, indicating that microglial metabolic reprogramming contributes to secondary inflammatory signaling beyond the lesion core.

Transient pharmacological inhibition of HK2 during the early post-injury window improved motor coordination despite extensive cortical damage, and partial genetic reduction of microglial HK2 phenocopied these benefits. These findings support two principles: early modulation of microglial metabolic programs can have durable functional consequences, and partial attenuation of HK2 activity is sufficient to improve outcome without abolishing essential microglial functions.

A key concern when targeting microglial metabolism is the risk of impairing beneficial processes, such as efferocytosis. In TREM2-deficient mice, failure of glycolytic reprogramming results in defective phagocytosis and worsened TBI pathology [6]. In contrast, HK2 antagonism in our model did not completely block activation. Instead, it reduced proliferation and inflammatory gene expression while preserving efferocytic capacity. Thus, partial HK2 inhibition appears to recalibrate inflammatory amplification rather than induce immune paralysis. This distinction is translationally important, as most TBI patients experience dysregulated inflammation rather than genetic immune deficiency.

The behavioral findings are consistent with this balanced model. HK2 antagonism improved motor coordination without affecting locomotion, anxiety-like behavior, or spatial working memory. The preservation of baseline cognitive performance argues against global neural or immune suppression and supports a selective effect on injury-induced dysfunction. Reduced inflammatory gene expression and ASC accumulation, particularly in the hippocampus, provide a mechanistic link between dampened inflammasome signaling and functional recovery. Excessive microgliosis and inflammasome activation contribute to synaptic dysfunction after TBI [22], and partial restoration of synaptophysin supports a model in which restrained inflammatory amplification stabilizes presynaptic integrity.

A notable feature of this study is the spatially restricted regulation of inflammasome activation within the hippocampal hilus. Although ASC protein expression was prominent at the cortical injury site, a similarly robust induction was observed in hilar microglia. The hilus is recognized as vulnerable to TBI-induced synaptic remodeling, interneuron loss, and altered neurogenesis, and plays a central role in regulating dentate gyrus excitability through mossy fiber circuitry [12–15]. Importantly, IL-1β signaling in this region suppresses neurogenesis and promotes circuit dysfunction [23]. The near-complete normalization of ASC levels in hilar microglia following HK2 antagonism, therefore, suggests that metabolic modulation of microglia may protect a functionally critical hippocampal subregion from sustained inflammatory amplification. Although neuronal and circuit-level consequences were not directly assessed, the spatial specificity of inflammasome suppression provides a strong rationale for future studies examining hippocampal network preservation.

The comparison between pharmacological and genetic targeting of HK2 further highlights important mechanistic nuances. Whereas LND treatment produced male-biased behavioral and transcriptional effects, partial genetic reduction of HK2 improved outcomes in both sexes. This dissociation indicates that the sex bias observed with LND is unlikely to reflect intrinsic HK2 biology, but rather sex-dependent pharmacokinetic or pharmacodynamic factors. Microglia exhibit sex-dependent responsiveness to systemic perturbations; antibiotic-mediated microbiome depletion induces male-biased microglial remodeling in development and AD [24,25], and metabolic stress induced by antiglycolytic agent, 2-deoxy-D-glucose elicits sexually dimorphic immune responses [26]. In this context, the male-biased effects of LND likely reflect compound-specific interactions within a sexually dimorphic immune-metabolic landscape rather than sex-restricted roles of HK2 itself.

Some limitations of the present study warrant consideration. The molecular mechanisms linking HK2 activity to inflammasome regulation require further definition, particularly given HK2’s non-canonical roles in mitochondrial tethering and inflammatory signaling [11,20,27–30]. Neuronal and circuit-level analyses were correlative, and direct electrophysiological measurements will be needed to establish causal links between reduced inflammasome activation and behavioral recovery. Additionally, although the genetic model isolates microglial HK2, pharmacological inhibition may influence other myeloid populations during later stages of injury.

In summary, our findings identify microglial HK2 as a modulable regulator of inflammatory amplification after severe TBI. Partial antagonism attenuates inflammasome activation, preserves efferocytic capacity, and improves motor recovery, while genetic reduction confirms the specificity of this effect. By recalibrating rather than abolishing microglial activation, HK2 targeting achieves a balance between repair and restraint that may better reflect physiological immune regulation. These findings position HK2 as a promising therapeutic node for limiting secondary inflammatory injury and preserving functional outcome after CNS trauma.

## Material and methods

### Animals

This study involved adult male and female C57BL/6J mice (16 weeks old) housed in the Indiana University School of Medicine (IUSM) animal care facility. Mice were maintained in accordance with USDA standards under a 12-h light/dark cycle with ad libitum access to food and water, following the Guide for the Care and Use of Laboratory Animals (National Institutes of Health, Bethesda, MD). All procedures were approved by the Indiana University Institutional Animal Care and Use Committee (IACUC) and were conducted in compliance with NIH guidelines.

We used two complementary strategies to investigate the role of hexokinase 2 (HK2) in traumatic brain injury. In the pharmacological approach, wild-type C57BL/6J mice were treated with Lonidamide, an HK2 inhibitor, after controlled cortical impact. In the genetic approach, C57BL/6J: CX3CR1-creERT2:HK2 flox/wt mice (called B6:HK2^Fl/wt^) received tamoxifen at 2 months of age to partially delete HK2, following the method described by Codocedo et al. [11]. These methods enabled us to examine both immediate pharmacological inhibition and a genetically induced reduction of HK2 in microglial activation after TBI. For the pharmacological part, mice were randomly assigned to four groups: (1) Sham + Vehicle, (2) Sham + Lonidamide, (3) controlled cortical impact (CCI) + Vehicle, (4) CCI + Lonidamide. For the genetic part, mice were randomly assigned to four groups: (1) Sham + oil, (2) Sham + Lonidamide, (3) CCI + oil, (4) CCI + tamoxifen. Both male and female mice participated in all experiments.

### Cell lines

The murine microglial cell line BV2 (RRID: CVCL_0182) was used as the efferocytic cell population and obtained from ATCC (Manassas, VA, USA; CRL-2467). BV2 cells were maintained in Dulbecco’s Modified Eagle Medium (DMEM) supplemented with 10% fetal bovine serum (FBS) and 1% penicillin–streptomycin.

The human neuroblastoma cell line SH-SY5Y (RRID:CVCL_0019, ATCC CRL-2266) was used as the source of dead-cell targets in efferocytosis assays. SH-SY5Y cells were cultured in DMEM/F12 medium supplemented with 10% FBS and 1% penicillin–streptomycin.

Both cell lines were maintained at 37 °C in a humidified incubator with 5% CO₂, used at low passage numbers, and routinely monitored for morphology and viability before experimental use.

### Drug Treatments

Lonidamine (LND) was purchased from Cayman Chemicals (14640), and a stock solution was prepared in DMSO (Sigma D2650). The daily dose of LND was 50 mg/kg body weight, diluted in PBS1x (DMSO 0.05%) and injected intraperitoneally into 4-month-old C57BL/6J mice at a volume of 1 mL for 7 consecutive days. The drug and vehicle solutions were well tolerated and did not affect the mice’s body weight. For in vitro experiments, 200 mM of LND was added to the media.

Tamoxifen (TAM) was obtained from Sigma (T5648). Stock solutions were prepared in vehicle (10% ethanol (Sigma E7023) + 90% corn oil (Sigma C8267)) and stored in the dark at −20°C. The daily dose of TAM was 15 mg/kg body weight, administered intraperitoneally to 2-month-old B6: CX3CR1-creERT2:HK2flox/wt mice in a volume of 0.1 mL for 5 consecutive days.

### Controlled Cortical Impact (CCI)

Mice were anesthetized with 5% isoflurane and secured in a stereotaxic frame, where anesthesia was maintained at 2% isoflurane. Meloxicam (5 mg/kg, subcutaneously) was administered for pain relief. The head was shaved, and a midline incision was made to expose the skull. A 4-mm craniotomy was performed on the left parietal bone between the lambda and bregma sutures, near the sagittal suture, using a rotary drill bit to preserve the dura. The impact site was at −1.7 mm posterior to the Bregma point, as defined by the mouse brain atlas of Franklin and Paxinos (Third Edition). A precision cortical impactor (Leica 3946393) was calibrated to zero at the dura. Care was taken to preserve the dura mater during surgery, as the impact device uses electrical conduction to determine the plane of the dura. The CCI was induced at a velocity of 3.0 m/s, with a dwell time of 00.1 seconds and an impact depth of 1.0 mm. Body temperature was maintained at 37°C throughout the procedure using a temperature-controlled heating pad.

Immediately after impact, the site was irrigated with sterile 0.9% sodium chloride to promote hemostasis. Surgi Foam was applied to absorb excess blood while avoiding deep pressure on the exposed tissue. Once hemostasis was confirmed, two drops of bupivacaine were applied to the impact site, and the wound was allowed to dry before closure. The incision was closed with tissue adhesive, carefully aligning the wound edges without disturbing the impact zone. Postoperatively, mice were placed in a temperature-controlled recovery incubator until fully awake and responsive (about 30 minutes), then returned to their home cages for ongoing monitoring. Mice were sacrificed 15 days after undergoing CCI. Mice were anesthetized with 1.2% 2,2,2-tribromoethanol (Avertin) and perfused with ice-cold phosphate-buffered saline (PBS).

### Behavioral Analysis

The mice were divided into four experimental groups for each of the previously described experimental strategies: pharmacological and genetic. The same mice tested on rotarod, spontaneous alternation, and Y maze were euthanized 15 days post-TBI and underwent brain perfusion for histopathology analysis.

#### Rotarod Test

To evaluate motor coordination and balance following traumatic brain injury (TBI), mice underwent the Rotarod test on a motorized rotating rod apparatus (Omnitech AccuRotor EzRod) with a 4 cm-diameter rod. The test was conducted 15 days after injury to monitor functional recovery. Mice were placed on a rotating rod set at 4 rpm, gradually increasing to 40 rpm over a specified duration. The latency to fall (the time in seconds before a mouse fell off the rod) was recorded for each trial. Each mouse completed three consecutive trials per session, with a rest period of a specified duration between trials to prevent fatigue. Mice received one day of training (3 trials at 4 rpm, walking for 60 seconds) 24 hours prior to the test. The average latency to fall was calculated for each session and compared between the TBI and control groups. A reduced latency to fall indicated motor impairment following TBI.

#### Y Maze test

The Y-maze test was used to assess spontaneous alternation behavior, an index of spatial working memory, following traumatic brain injury. The apparatus consisted of three identical white plastic arms (A, B, and C), each approximately 35 cm long and 4 cm wide, arranged at 120° angles. Testing was conducted under low-light conditions (∼5 lux) to minimize anxiety-related confounds. Mice were placed individually at the center of the maze and allowed to explore all three arms freely for 10 minutes. Arm entries and movement patterns were recorded and analyzed using ANY-Maze software (Stoelting Co). Spontaneous alternation (%) was calculated as the number of consecutive triads containing entries into all three arms without repetition, divided by the total number of possible alternations. Measurements were analyzed at 5, 8, and 10 minutes to assess working memory performance over time. A reduction in the percentage of spontaneous alternation was interpreted as impaired working memory function.

#### Open Field and Open Field Anxiety

General locomotor activity and exploratory behavior were assessed using the open field test. The apparatus consisted of a white plastic square arena measuring approximately 40 × 40 cm with 30 cm-high walls. Testing was conducted under low-light conditions (∼5 lux) to minimize stress-induced behavior. Mice were placed individually in the center of the arena and allowed to explore freely for the duration of the trial. Movements were recorded and analyzed using ANY-Maze software (Stoelting Co.).

Anxiety-like behavior was evaluated in the same open field arena by quantifying time spent in the center, periphery, and corner zones. Reduced exploration of the center, along with increased time spent in the periphery or corners, was interpreted as increased anxiety-like behavior. All analyses were conducted by an experimenter blinded to injury and treatment conditions.

### Tissue collection and homogenization

At the experimental endpoint, animals were anesthetized and euthanized in accordance with procedures approved by the Indiana University School of Medicine (IUSM) Institutional Animal Care and Use Committee. After transcardial perfusion with 1X PBS, mouse brains were quickly removed and separated into ipsilateral (injured) and contralateral hemispheres, which were then collected separately for further analyses.

For biochemical studies, both the ipsilateral and contralateral hemispheres were homogenized separately in tissue homogenization buffer (THB) supplemented with a protease inhibitor cocktail. One part of each homogenate was further sonicated to extract protein and used for Western blot analyses. The remaining part was stored at −80 °C in RNA-Bee (Amsbio, CS-501B) for later RNA extraction and gene expression analysis. For histopathological analysis, whole brains were collected after perfusion, fixed overnight in 4% paraformaldehyde at 4 °C, cryoprotected in 30% sucrose at 4 °C, embedded, and then processed on a cryostat into 30 µm free-floating sections.

### Western blot

After sonication, the soluble fraction of brain lysates was obtained by centrifugation at 14,000 rpm for 12 minutes. Protein concentrations were measured using the Pierce BCA assay (Thermo-Fisher, 23225). For protein identification and relative quantification, 5 to 20 μg of protein was loaded onto Bolt 4%–12% Bis-Tris Plus Protein gels (Thermo-Fisher, NW04122BOX) and run at 200 mV for 40 minutes. Proteins were then transferred to a PVDF membrane (Millipore, IPVH00010) at 400 mA for 90 minutes at 4 °C. After one hour of blocking in 5% BSA (in PBS 1X), primary antibodies were applied overnight in blocking buffer at 4 °C: HK2 (1:1000, Abcam ab209847), HK1 (1:1000, Santa Cruz sc-46695), Trem2 (1:1000, R&D Systems AF1729), ASC (1:1000, Cell Signaling 67824S), and Beta Actin (Santa Cruz sc-130065). HRP-conjugated secondary antibodies were incubated for one hour at room temperature at the appropriate dilutions. The signal was developed using Immobilon Western Chemiluminescent HRP Substrate (Millipore, WBKLS0500). Images were captured with an Invitrogen iBright 1500 Imagers (Thermo Fisher Scientific), and protein quantification was performed by measuring the optical density of the specific bands using ImageJ (NIH). Samples from all experimental groups were loaded in each experiment and normalized to their respective controls. For statistical analysis, the normalized data from independent experiments were combined.

### RNA isolation and quantitative real-time PCR

RNA was extracted from homogenized tissue using the PureLink RNA Mini Kit (Invitrogen, Ref 12183025) following the manufacturer’s instructions. The RNA was then reverse-transcribed into cDNA with the High-Capacity RNA-to-cDNA Kit (ThermoFisher Scientific, 4387406). TaqMan Gene Expression MasterMix (Applied Biosystems, 4369016) was used for qPCR according to the manufacturer’s instructions. For all mRNA analyses, the housekeeping gene GAPDH (Mouse GAPDH, Thermo Fisher Scientific, 4352339E) was employed. Results are expressed as the relative fold change in gene expression, normalized to the wild-type calibrator. For statistical analysis, the relative delta Ct method was used.

### Immunostaining, image acquisition, and image analysis

Immunohistochemical analyses were performed on coronal brain sections spanning the rostrocaudal extent of the injury, from approximately bregma −0.82 to bregma −2.92. These coordinates served as anatomical references and may vary slightly between animals. Representative sections were selected from each region, with particular emphasis on the impact zone, defined as the pericontusional area between bregma −1.70 and −2.46, where the primary injury was induced. At least five free-floating sections per brain were analyzed to ensure adequate regional representation.

Free-floating sections were washed and permeabilized in 0.1% Triton in PBS (PBST), and antigen retrieval was performed using 1X Reveal decloacker (Biocare Medical) at 85 °C for 15 min. Sections were blocked in 5% normal donkey serum in PBST for one hour at room temperature (RT). Primary antibodies were incubated in 5% normal donkey serum in PBST overnight at 4 °C. Sections were immunostained with antibodies against HK2 (ab209847), Iba-1 (microglial marker, NB100-1028), and ASC (inflammasome adaptor protein, Cell Signaling 67824S). All sections were processed in parallel under identical conditions to minimize technical variability. Sections were washed and visualized using species-specific AlexaFluor fluorescent antibodies (diluted 1:1000 in 5% normal donkey serum in PBST for one hour at RT). Sections were counterstained and mounted onto slides.

Immunostained brain sections were imaged with a Leica DM6 B inverted microscope at 10X magnification to assess lesion morphology across defined anatomical regions. To evaluate immunoreactive areas and lesion extent, images were acquired at 10X magnification and a 1 µm z-step, enabling systematic sampling through the tissue thickness and quantitative comparison between ipsilateral and contralateral hemispheres across the rostrocaudal extent of the injury. HK2 and ASC expression within microglia in the hippocampal region was assessed at the same 10X magnification. Higher-resolution imaging of peri-contusional cortical tissue was performed at 20X magnification. All imaging parameters were held constant across experimental groups to ensure reproducibility of quantitative analyses.

### Efferocytosis Assay

Efferocytosis assays were performed using BV2 microglial cells as the phagocytic population and fixed SH-SY5Y neuroblastoma cells as apoptotic targets, following a previously validated quantitative protocol with minor adaptations [31,32]. Briefly, SH-SY5Y cells were incubated with 5-chloromethylfluorescein diacetate (CellTracker Green CMFDA dye, Invitrogen, US) diluted in DMEM to a 20 μM final concentration for 20 min at 37°C, followed by two washes with 1X PBS. Subsequently, cells were dissociated with trypsin and fixed with paraformaldehyde to render them nonviable. Labeled SH-SY5Y cells were resuspended in DMEM and added to BV2 cells at a defined 1:1 cell ratio. Co-cultures were maintained under standard culture conditions, and efferocytosis was monitored by live-cell imaging with an Omni imaging system (Axion BioSystems, Inc., USA). Internalization of SH-SY5Y apoptotic cells was identified by the decrease in fluorescent objects, indicating successful engulfment and degradation.

## QUANTIFICATION AND STATISTICAL ANALYSIS

Statistical tests and the number of animals analyzed are indicated in the figure legends. Comparisons between two groups were conducted with unpaired Student’s t-tests or paired t-test when comparing changes between the contralateral vs ipsilateral side of the injury. Comparisons among multiple groups were conducted using one-way ANOVA with Tukey’s post hoc multiple comparisons test in Prism 10 (GraphPad, San Diego, CA, United States). Error bars represent the standard error of the mean (SEM). Statistical significance was met when the p-value was less than 0.05.

## Acknowledgment

This work was supported by the Behavioral Phenotyping Core (BPC) at the Indiana University School of Medicine.

## Author contributions

Conceptualization: CMR and JFC. Methodology: CMR and JFC Investigation: CMR, JFC, PBF, and JS. Formal analysis: CMR, JFC and PBF. Writing, review, and editing: CMR, JFC, and GEL. Funding Acquisition CMR, JFC, and GEL. All authors have read and agreed to the published version of the manuscript.

## Availability of Data and Materials

No publicly available data or shared data are cited. All data needed to evaluate the conclusion of the current study are present in the paper and/or the supplementary materials. Additional data are available from the corresponding author on request.

---

**Supplementary Figure 1.**
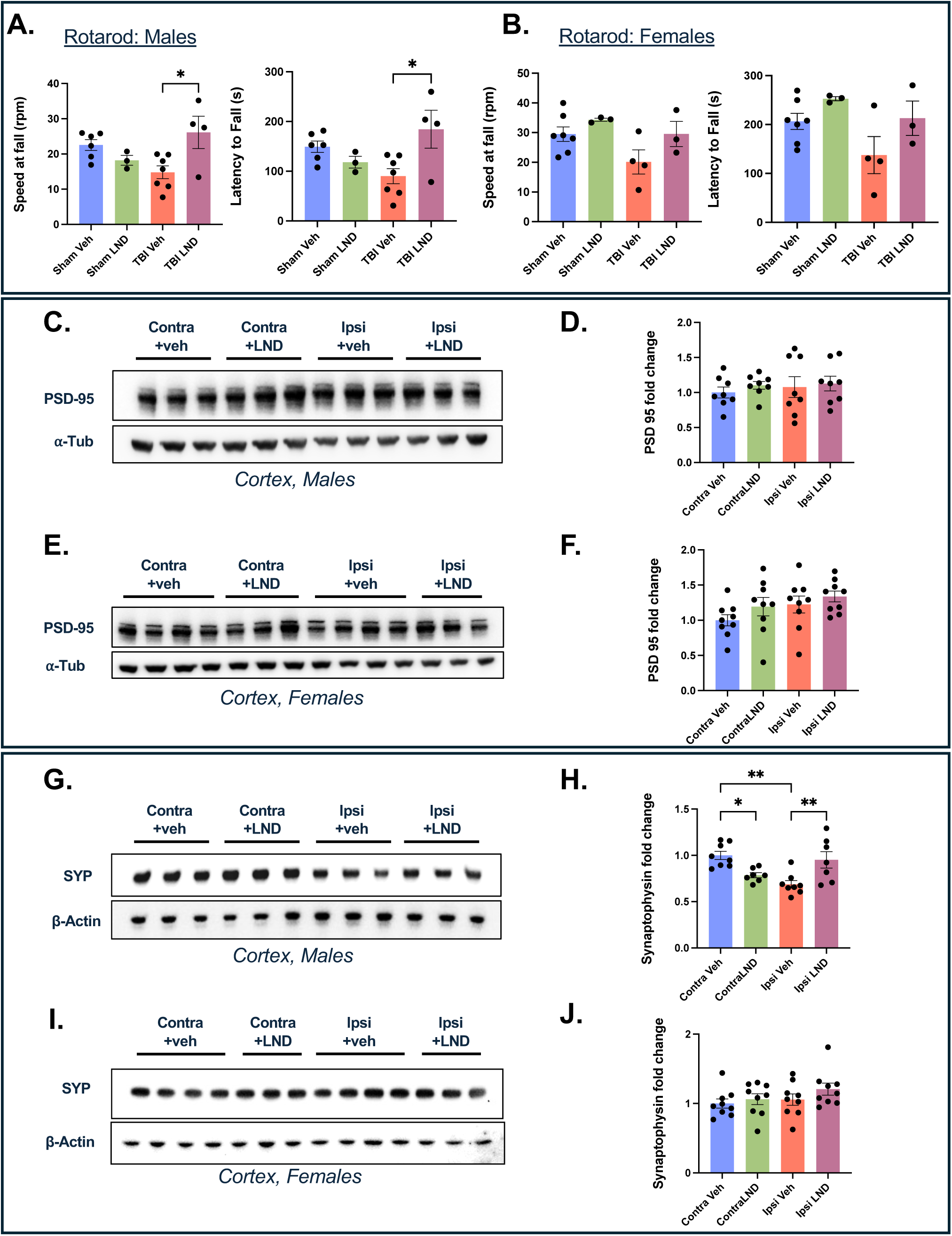
Sex-dependent effects of HK2 antagonism on motor behavior and synaptic protein expression. **(A–B)** Rotarod performance was analyzed separately in male and female mice following vehicle or LND treatment. Behavioral improvement is primarily observed in male mice (*n* = 4-7 for males and n=3-7 for females per group, *p < 0.05, Ordinary one-way ANOVA with post hoc Tukey’s multiple comparisons). **(C–F)** Western blot analysis and quantification of PSD-95 in cortical tissue show no significant effects of unilateral CCI or LND treatment. **(G–H)** Synaptophysin (SYP) levels are reduced in the male cortex following CCI and partially restored by LND treatment. (*n* = 7-8 per group, *p < 0.05 and *p < 0.01, Ordinary one-way ANOVA with post hoc Tukey’s multiple comparisons) **(I–J)** SYP expression in the female cortex is not significantly altered by LND.

**Supplementary Figure 2.**
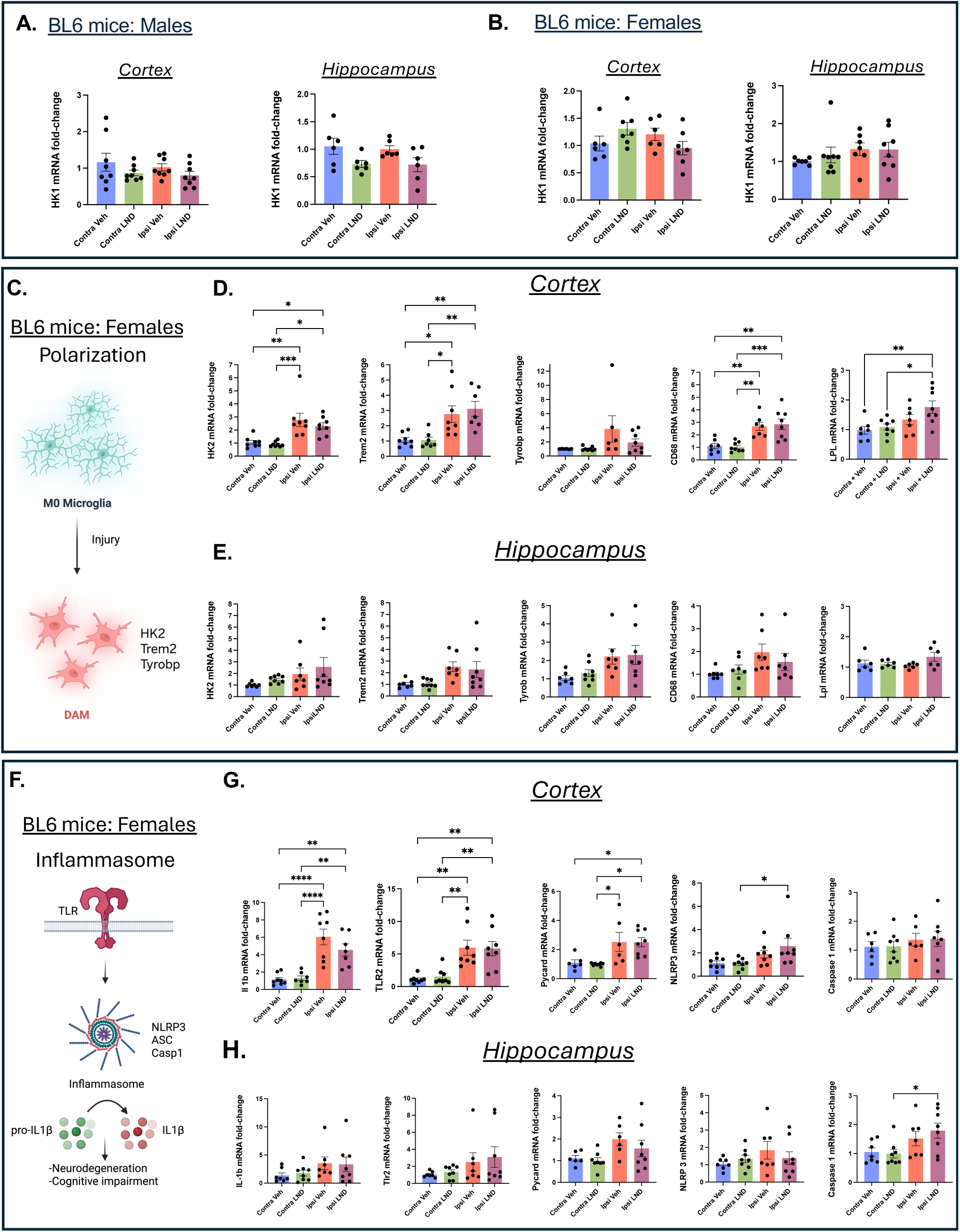
HK2 antagonism does not significantly alter microglial gene expression in female mice after TBI. **(A)** qPCR analysis of HK1 in the cortex and hippocampus of male mice following CCI and LND treatment. **(B)** qPCR analysis of HK1 in the cortex and hippocampus of female mice following CCI and LND treatment. **(C-E)** Expression of microglial activation–associated genes in the cortex and hippocampus of female mice following CCI and LND treatment. (n = 6-8 per group, *p < 0.05, **p < 0.01, and ***p < 0.001, Ordinary one-way ANOVA with post hoc Tukey’s multiple comparisons) **(F-H)** Inflammasome-related gene expression in female mice shows no significant modulation by LND in either region. (*n* = 6-7 per group, *p < 0.05, **p < 0.01, and ***p < 0.001, Ordinary one-way ANOVA with post hoc Tukey’s multiple comparisons)

**Supplementary Figure 3.**
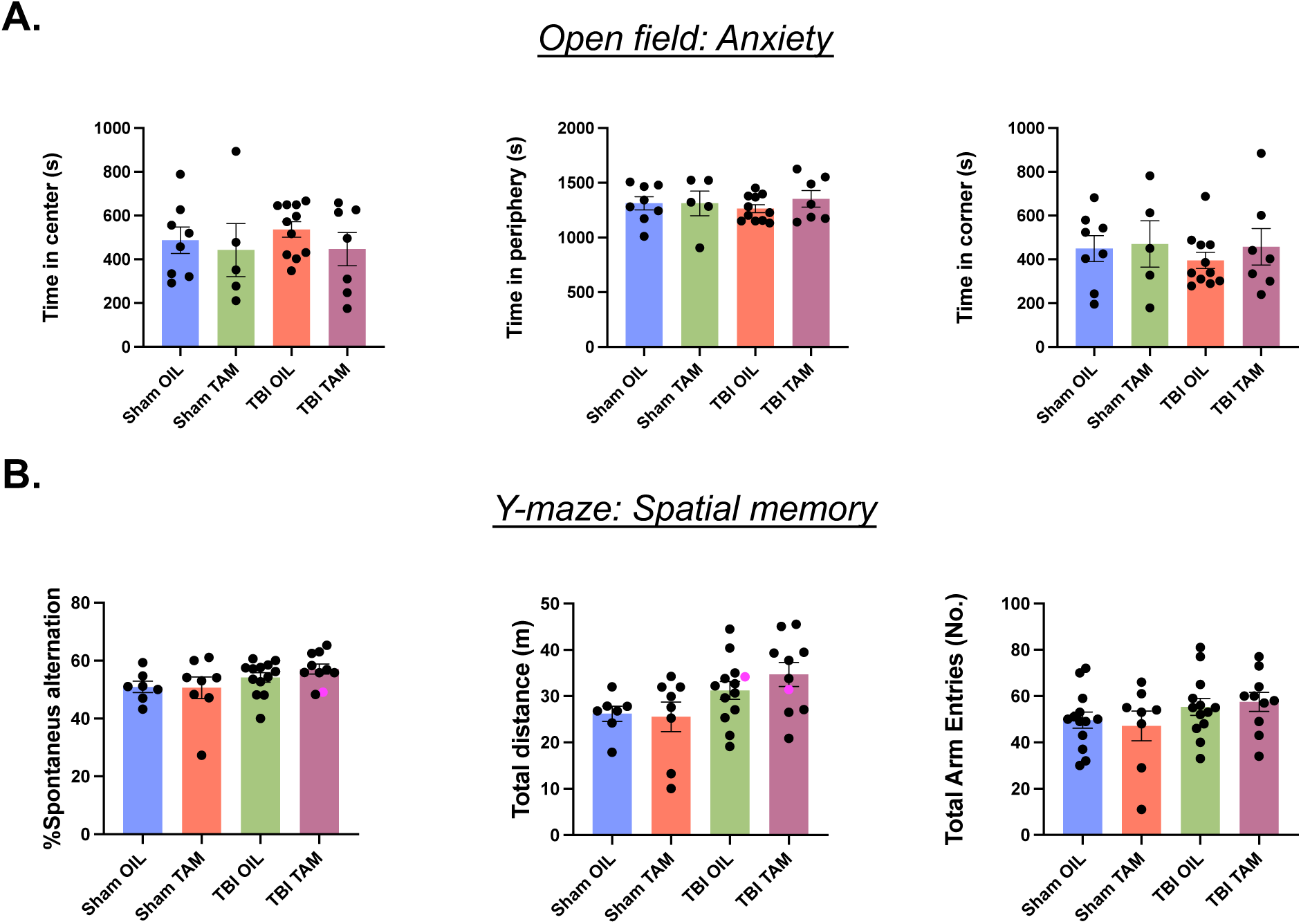
Partial genetic reduction of HK2 does not affect anxiety-like behavior or spatial memory. **(A)** Open field center exploration. **(B)** Y-maze spontaneous alternation.

